# CardioClassifier – demonstrating the power of disease- and gene-specific computational decision support for clinical genome interpretation

**DOI:** 10.1101/180109

**Authors:** Nicola Whiffin, Roddy Walsh, Risha Govind, Matthew Edwards, Mian Ahmad, Xiaolei Zhang, Upasana Tayal, Rachel Buchan, William Midwinter, Alicja E Wilk, Hanna Najgebauer, Catherine Francis, Sam Wilkinson, Thomas Monk, Laura Brett, Declan P O'Regan, Sanjay K Prasad, Deborah J Morris-Rosendahl, Paul JR Barton, Elizabeth Edwards, James S Ware, Stuart A Cook

## Abstract

**Purpose:** Internationally-adopted variant interpretation guidelines from the American College of Medical Genetics and Genomics (ACMG) are generic and require disease-specific refinement. Here we developed CardioClassifier (www.cardioclassifier.org), a semi-automated decision-support tool for inherited cardiac conditions (ICCs).

**Methods:** CardioClassifier integrates data retrieved from multiple sources with user-input case-specific information, through an interactive interface, to support varian interpretation. Combining disease- and gene-specific knowledge with variant observations in large cohorts of cases and controls, we refined 14 computational ACMG criteria and created three ICC-specific rules.

**Results:** We benchmarked CardioClassifier on 57 expertly-curated variants and show full retrieval of all computational data, concordantly activating 87.3% of rules. A generic annotation tool identified fewer than half as many clinically-actionable variants (64/219 vs 156/219, Fisher’s **P**=1.1x10-18), with important false positives; illustrating the critical importance of disease and gene-specific annotations. CardioClassifier identified putatively disease-causing variants in 33.7% of 327 cardiomyopathy cases, comparable with leading ICC laboratories. Through addition of manually-curated data, variants found in over 40% of cardiomyopathy cases are fully annotated, without requiring additional user-input data.

**Conclusion:** CardioClassifier is an ICC-specific decision-support tool that integrates expertly curated computational annotations with case-specific data to generate fast, reproducible and interactive variant pathogenicity reports, according to best practice guidelines.

## INTRODUCTION

Inherited cardiac conditions (ICCs) represent a major health burden with a combined prevalence of ~1%^1^. Genetic testing is recommended to support the management of many ICCs, with roles in diagnosis (particularly valuable for identification of at-risk relatives), prognostication, and therapeutic stratification. The principle challenge in genetic testing across all diseases is the interpretation of identified sequence variants. This requires evaluation of data from diverse sources, including clinical observations, computational data and data derived from the literature. Although existing tools aid collection of some of these data types, scientists and clinicians must often access multiple data sources whilst interpreting a single genetic variant.

The American College of Medical Genetics and Genomics (ACMG) and the Association for Molecular Pathology (AMP) recently released guidelines that aim to standardise variant interpretation^2^. These guidelines outline a set of evidence criteria to assess each variant against, along with how these might be weighted and combined to reach a classification. Studies have, however, shown that even when following the ACMG/AMP guidelines, interpretation can differ between different laboratories, with discordance in excess of 10%^3^. One key reason for this discordance is the structure of the ACMG/AMP guidelines, which are deliberately broad and lack specific thresholds, to allow adoption across the full spectrum of genetic disorders. As a result, the challenge to individual disease domains is to incorporate expert gene and disease-specific knowledge, to optimise variant interpretation and introduce consensus. Initiatives such as the Clinical Genome Resource (ClinGen)^4^, are working to define such disease- and gene-specific thresholds, although these are currently limited to pilot phases for specific gene-disease pairs.

The introduction of guidelines, including the logic behind reaching each classification, opens the way for new computational solutions to facilitate their adoption and increase consistency. Indeed, publication of the guidelines has led to the emergence of interactive tools^5,6^, however, to date only one of these builds in automation^7^, and none incorporate expert disease-specific knowledge.

Here, we describe CardioClassifier, a powerful new tool that utilises the framework outlined by the ACMG/AMP guidelines, to automatically annotate variants across 17 computational criteria. Each criterion has been individually parametrised for each gene-disease pair using expert disease-specific knowledge. Automated data are integrated with interactively added case-specific information to calculate variant pathogenicity in a fully interactive web-interface that represents a comprehensive variant interpretation platform for ICCs.

## MATERIALS AND METHODS

We have initially implemented CardioClassifier for a set of 40 genes with definitive links to 11 ICCs. The full list of genes and corresponding disease phenotypes can be found in Supplementary Table 1. The following section describes the development and optimisation of CardioClassifier and is split into three main sections

1. Rule selection and optimisation – describes how the individual ACMG/AMP criteria were adapted and parameterised for ICCs
2. Code and implementation – outlines the technical details of the tool
3. Benchmarking – describes the data and process used to fully test CardioClassifier

### Rule selection and optimisation

For each rule in the ACMG/AMP framework, we first evaluated whether the rule was applicable to the ICC under investigation and, where appropriate, defined more precisely the circumstances under which the rule would be activated. For example, we defined disease-specific allele frequency thresholds taking into account the genetic architecture of each disease^8^, to dictate activation of BS1 (minor allele frequency is too common for the disorder), and used genetic burden data^9^ along with extensive literature curation to determine for which genes and diseases loss-of-function mutations are a disease mechanism (activation of PVS1). As part of this development process, we compared rule activation in CardioClassifier to a set of variants manually curated as part of routine clinical service at the Royal Brompton Hospital (see Supplement). Full details of how each rule is parameterised can be found in the Supplement to this manuscript.

In addition to the 14 rules outlined in the ACMG/AMP guidelines, we have created three modified rules that enhance the interpretation of ICC variants. These rules are fully described in the results section and in the Supplement.

### Code and implementation

CardioClassifier is implemented server-side in perl and PHP. Users upload variant data in the form of a single sample variant call file (VCF) or individual variant details, which are then annotated by the Ensembl variant effect predictor (VEP)^10^ and converted to a table using the tableize_vcf.py script within LOFTEE (https://github.com/konradjk/loftee). Protein altering and splice site variants (coding ±8bps) are analysed for a set of 40 genes associated with inherited cardiac conditions (Table 1). We look to continuously expand this list, focusing on curated genes robustly implicated in disease, emerging from community efforts such as ClinGen^4^.

**Table 1:** Details of gene-disease pairs currently analysed by CardioClassifier. The disease class column details the larger sub-panels relating to broad disorder types that each disease and gene set are within.

| Disease | Disease Class | Genes | Total Genes |
| --- | --- | --- | --- |
| DCM | Cardiomyopathy | LMNA, TNNT2, SCN5A, TTN, TCAP, MYH7, VCL, TPM1, TNNC1, RBM20, DSP, BAG3 | 12 |
| HCM | Cardiomyopathy | MYH7, TNNT2, TPM1, MYBPC3, PRKAG2, TNNI3, MYL3, MYL2, ACTC1, CSRP3, PLN, TNNC1, GLA, FHL1, LAMP2, GAA | 16 |
| ARVD/C | Cardiomyopathy | DSP, PKP2, DSG2, DSC2, JUP | 5 |
| RCM | Cardiomyopathy | TNNI3 | 1 |
| ncCM | Cardiomyopathy | MYBPC3, MYH7 | 2 |
| Noonan syndrome | Cardiomyopathy | RAF1, SOS1, PTPN11, KRAS | 4 |
| Long QT syndrome | Arrhythmia | KCNQ1, KCNH2, SCN5A, KCNE1, KCNE2 | 5 |
| Brugada syndrome | Arrhythmia | SCN5A | 1 |
| CPVT | Arrhythmia | RYR2 | 1 |
| Marfan syndrome | Aortopathy | FBN1 | 1 |
| FH | - | LDLR | 1 |

The classifier automatically assesses each variant for 17 rules across three distinct data categories, as defined by the ACMG/AMP guidelines^2^. It also consults an internal knowledge base of additional evidence, grouped by ACMG rule, either derived from community curation efforts or manually curated internally. The output is displayed on a PHP webpage that allows the user to interact and add (or remove) additional levels of evidence.

### Benchmarking

#### Datasets

In order to test CardioClassifier extensively we used data from the following sources:

1. ClinVar – all variants identified as ‘Pathogenic’ or ‘Likely Pathogenic’ by multiple submitters with no conflicting data (i.e. reports of ‘Benign’ or ‘Likely Benign’ or ‘Uncertain Significance’) for hypertrophic cardiomyopathy (HCM; n=158), dilated cardiomyopathy (DCM; n=16), long QT syndrome (LQTS; n=18), catecholaminergic polymorphic ventricular tachycardia (CPVT; n=1), Brugada syndrome (Brs; n=4) or arrhythmogenic right ventricular cardiomyopathy (ARVC; n=22) were extracted from the 20161201 release of ClinVar^11^ using publically available scripts^12^.
2. ClinVar – a wider set of missense variants clinically curated (i.e. not literature only or research) as ‘Pathogenic’ or ‘Likely Pathogenic’ for LQTS from one or more submitter with at least one review status star (n=48).
3. 57 protein-altering variants that have been expertly curated as part of the recent ClinGen pilot for **MYH7**.
4. A prospective dataset of 327 HCM cases and 625 healthy volunteers recruited to the NIHR Royal Brompton cardiovascular BRU, all phenotypically characterised using cardiac MRI. Samples were sequenced using the IlluminaTruSight Cardio Sequencing Kit^1^ on the Illumina NextSeq platform. This study had ethical approval (REC: 09/H0504/104+5) and informed consent was obtained for all subjects.

#### Comparison with existing resources

Although no similar tools currently exist that are both automated and disease-specific, in February 2017 Li and Wang released InterVar^7^ as the first tool to build in a level of automation. We compared the performance of CardioClassifier against InterVar to assess the importance of our disease-specific annotations. We used the ClinVar dataset of 219 variants described above as a test dataset.

InterVar scripts were downloaded from GitHub (https://github.com/WGLab/InterVar) and individually run for each disease using an engineered VCF file. To ensure a fair comparison, we edited the ‘disorder_cutoff’ to be equivalent to the thresholds used to activate BS1 in CardioClassifier. All other settings were left as default and no additional evidence was uploaded. We compared both the final classifications and the individual rules that were activated by each tool.

#### Coverage comparison between ExAC and healthy volunteers

Variants that are common in the general population are unlikely to be disease-causing. CardioClassifier therefore filters variants relative to the exome aggregation consortium (ExAC) dataset of 60,706 indivduals^13^. As this dataset is derived from exome sequencing data, some genomic regions, especially those that are repetitive or with high GC content, are not fully covered in these data. To overcome this limitation, we have included an additional cohort of 1383 healthy volunteers (625 UK and 758 from Singapore) sequenced on the TruSight Cardio sequencing panel, which has been optimised to achieve near complete coverage across 174 genes involved in ICCs^1^. In addition, this dataset identifies Illumina platform-specific technical artefacts, allowing us to distinguish true differences in case and control frequencies from sequencing errors.

To assess the contribution of our healthy volunteer cohort to variant frequency filtering we compared the coverage in these samples to the ExAC dataset (release 0.3.1) across the coding regions of the 40 genes currently parameterised within CardioClassifier (total 0.25 Mb). We report metrics as ‘sample bases’ where each base in each sample is individually assessed for coverage >20x, and the proportion of these meeting this threshold are reported.

## RESULTS

### Semi-automation leads to high quality and reproducible variant interpretation

CardioClassifier provides a simple-to-use web interface that takes as input either individual variant details or a single sample VCF (Supplementary Figure 1). Where a diagnosis is known, users select one of 11 cardiac disorders for which to perform analysis that pre-specifies disease gene selection by CardioClassifier. Where a diagnosis is more uncertain (e.g. sudden cardiac death or complex cardiomyopathy), analysis can be performed over all 40 genes currently parameterised within the tool, or a subset associated with related disorders (e.g. all cardiomyopathies, or all arrhythmia syndromes; Table 1). Details of the key features of CardioClassifier can be found in Table 2.

**Table 2:** Key features of CardioClassifier. Included are details of each key feature and which of three currently available tools (Alamut, InterVar and the ClinGen pathogenicity calculator) also includes each feature.

| Feature | Description | CardioClassifier | Alamut | InterVar | ClinGen Pathogenicity Calculator |
| --- | --- | --- | --- | --- | --- |
| Collates data from multiple sources | CardioClassifier retrieves data from multiple databases/resources including ExAC, ClinVar, ACGV and dbNSFP as well as internally derived data | ✓ | ✓ | ✓ | - |
| Takes a standard VCF or variant details as input and annotates with effect on sequence and protein | The Ensembl Variant Effect Predictor is used to annotate all variants according to protein consequence | ✓ | ✓ | ✓ | - |
| ACMG/AMP rules parameterised through expert curation according to specific gene and disease | We have developed expertly-curated gene and disease specific thresholds for 14 computational ACMG/AMP criteria in addition to 3 specifically created ICC specific rules. | ✓ | - | - | - |
| Computational data used to activate ACMG/AMP rules | Each variant is automatically assessed against 17 computational criteria | ✓ | - | ✓ | - |
| Interactive refinement of rules and addition of case-level data | Users can interactively add or remove evidence pertaining to any of the ACMG/AMP rules | ✓ | - | ✓ | ✓ |
| Integration of automated annotations and case-level interactive additions to calculate a classification according to the ACMG logic | The logic from the ACMG/AMP guidelines is used to provide a final classification | ✓ | - | ✓ | ✓ |
| Evidence used to generate classification displayed | The thresholds and data used in CardioClassifier is transparent and printed on the report | ✓ | - | - | - |
| Knowledge base of case-level annotations | We have created a 'knowledge base' whereby manually curated case-level evidence is stored and used to populate variant reports | ✓ | - | - | - |

Each inputted variant is annotated for up to 17 computational criteria, with results outputted to a pre-populated grid representing the ACMG/AMP framework (Figure 1). Each criterion has been parameterised using expert disease and gene specific knowledge (see supplement for full details), leading to high quality and reproducible annotation. The variant report is interactive, allowing a user to add additional case-level evidence to generate and refine a final classification (Supplementary Figure 2). The report is transparent, with all evidence used to calculate the classification displayed, along with links out to eight external resources that are commonly used for interpretation of ICC variants: the ExAC browser^14^, Ensembl, the UCSC genome browser, ClinVar, PubMed, Google, the Beacon Network (https://beacon-network.org) and the Atlas of Cardiac Genetic Variation (ACGV)^9^.

**Figure 1:**
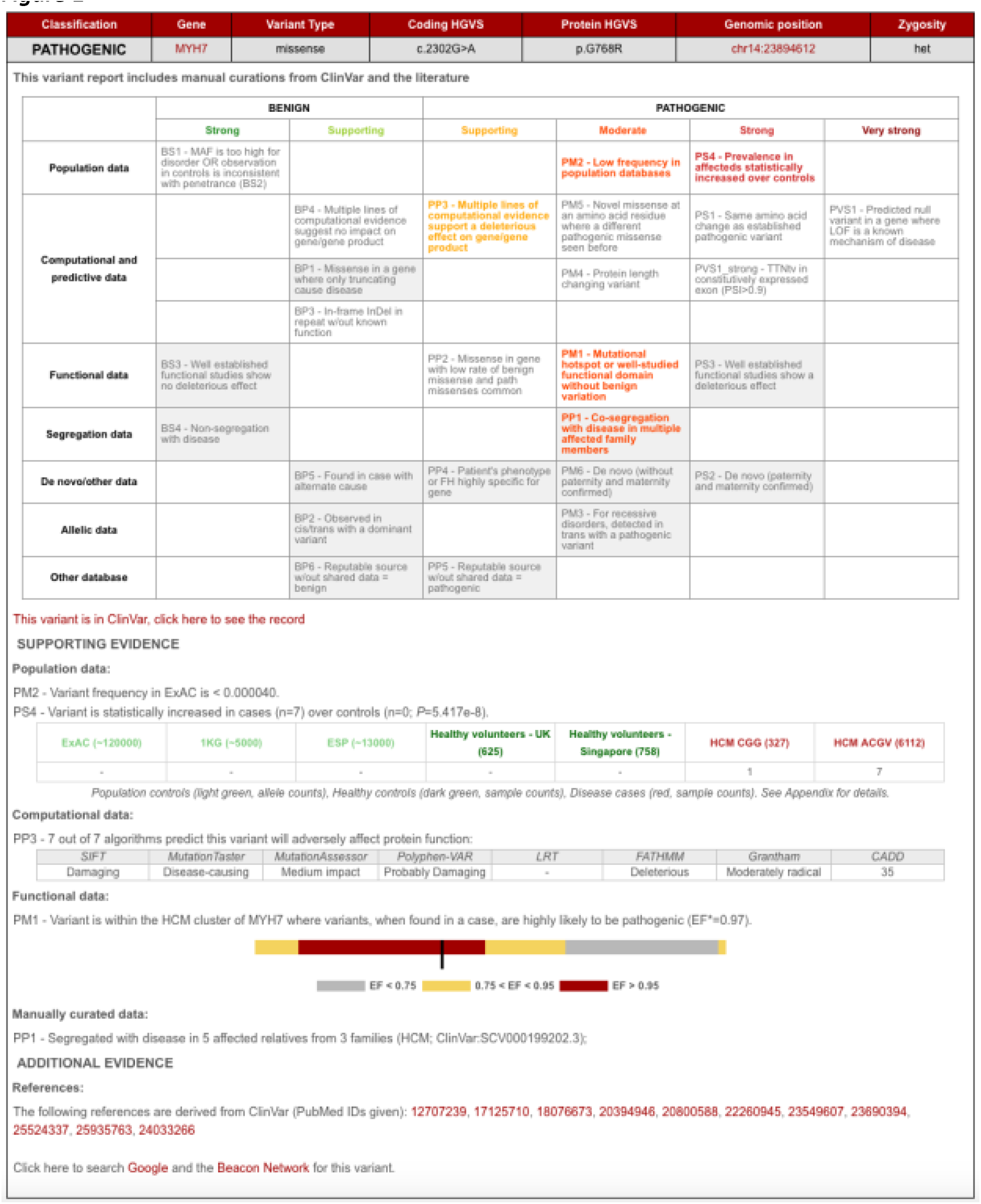
Example variant report output by CardioClassifier. A grid is output for each individual variant. Rules highlighted in colour are activated for the variant and rules in grey on a white background are assessed but not activated. A user can click on a rule to manually add or remove a piece of evidence. All evidence used to assess the variant is displayed under the grid along with links out to external resources. An overall classification for the variant using the ACMG/AMP logic is displayed in the top left corner. *EF – etiological fraction; the prior probability that a variant, identified in a case, is Pathogenic^9^.

### Disease-specific optimisation of ACMG/AMP rules

For seven computational criteria (PS1, PM4, PM5, PP3, BA1, BP3 and BP4), parameterisation is consistent across all gene-disease pairs, however, for the remaining criteria, we have incorporated expert disease, gene and variant-type specific knowledge and data to define thresholds for activation. This includes determination of robust disease-specific maximum frequency thresholds^8^ (BS1 and PM2; Supplementary Table 1), and using large disease cohorts to define both ‘mutational hotspots’^9^ (PM1; Figure 2a) and variants observed more frequently in cases when compared with population controls (PS4).

**Figure 2:**
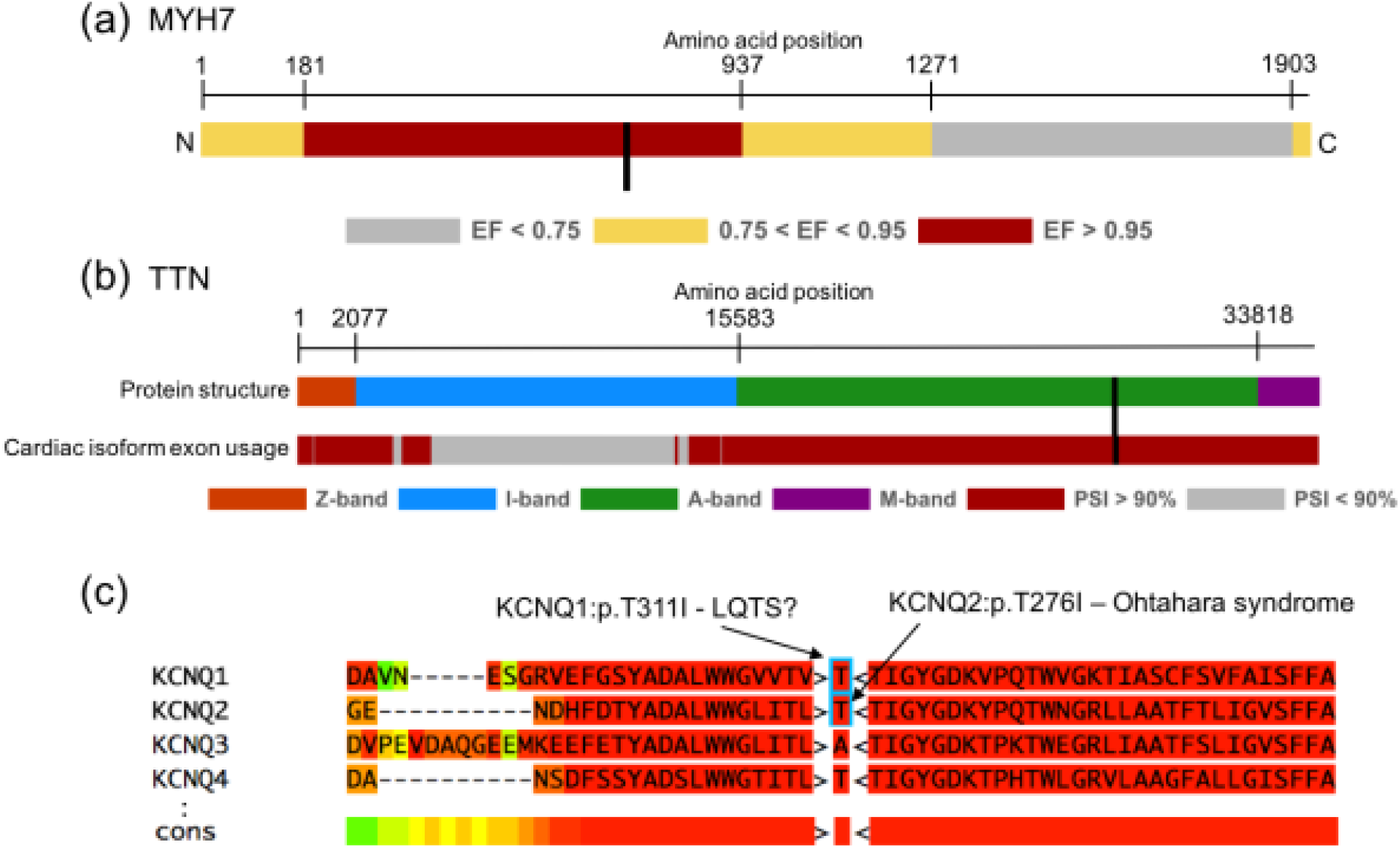
>Examples of disease-specific optimisation of ACMG/AMP rules. (a) Missense variants within a sub-portion of MYH7, when identified in a HCM patient, have a 97% prior probability of being Pathogenic (etiological fraction; EF=0.97). We activate PM1 for missense variants in this region. Here we use **MYH7**:c.2221G>T as an example (labelled with a black bar). (b) Truncating variants in **TTN** are only known to cause DCM when found in exons constitutively expressed in the heart (proportion spliced in (PSI) > 0.9). We activate PVS1_strong for these variants. Here we use **TTN**:c.86641delC as an example (labelled with a black bar). (c) Variants that have been identified as Pathogenic in paralogous genes may identify residues that are intolerant to variation. We have created two modified rules, PS1_moderate and PM5_supporting to incorporate this evidence. Here we use **KCNQ1**:p.T311I as an example. **KCNQ2**:p.T276I is associated with Ohtahara syndrome. We activate PS1_moderate for **KCNQ1**:p.T311I which is the equivalent missense change (i.e. same reference and alternate amino acids) in a different member of the same protein family.

As most large reference datasets, such as ExAC, have not been specifically screened for ICCs, it is expected that low frequency disease-associated alleles will be observed. This is particularly the cases for ICCs, which often manifest later in life and exhibit incomplete penetrance. We have therefore modified PM2 so as not to inappropriately discard variants seen at very low frequencies in these reference datasets.

In addition, we have created extensions to three ACMG/AMP rules, to enhance interpretation of ICC variants. Firstly, we have modified PVS1 for the titin (TTN) gene, which has a role in up to 20% of dilated cardiomyopathy (DCM) cases^15^. We have previously shown that only TTN truncating variants (TTNtv) in exons constitutively expressed in the heart are robustly associated with DCM^15^. Additionally, it is unclear that the mechanism of action for these variants is truly loss of function (LoF). Instead of scoring all TTNtv equally and assuming an underlying LoF mechanism, we only score TTNtv highly if they are in constitutive exons (proportion spliced in (PSI) > 0.9), and we reduce the strength of evidence by one level from very strong to strong (coded as PVS1_strong).

We also included modifications to PS1 and PM5 that utilise known disease-causing variants in related genes/proteins (paralogues), which likely identify residues intolerant to variation^16^ (Figure 2c). Where nothing is known about variants at the equivalent residue of the same gene, we use high confidence variants as evidence if they affect the equivalent residue in a paralogue (with the same reference allele), either with the same substitution (rule PS1_moderate – **Equivalent amino acid change as an established pathogenic variant in a paralogous gene**), or a different substitution (rule PM5_supporting – **Missense change at an amino acid residue where a pathogenic missense change has been seen in the equivalent residue of a paralogous gene**). This analysis is currently restricted to the families of predominantly ion channel proteins associated with inherited arrhythmia syndromes for which this method has been previously validated^16,17^.

We have previously shown paralogue annotation to be informative for over one third of novel SNVs^17^, and independent validation has shown a high specificity and PPV compared with other sources of evidence^18,19^. To determine the effect of these criteria on variant classification (before inclusion of any case-level or functional data that cannot be computationally predicted) we used 48 clinically curated missense variants from ClinVar identified as ‘Pathogenic’ or ‘Likely Pathogenic’ for LQTS. Paralogue annotation rules were activated for 11/48 (22.9%) variants and resulted in a change of class from variant of uncertain significance (VUS) to Likely Pathogenic for 63.6% of these (7/11).

### Highly curated datasets of disease cases and healthy controls aid annotation and filtering

As well as publically available data for both cases and population controls, CardioClassifier incorporates data from three highly-curated in-house datasets sequenced with the Illumina TruSight Cardio sequencing panel^1^. Counts from 877 DCM, 327 HCM cases, and 1383 healthy volunteers, all rigorously phenotyped using cardiac MRI, are used to annotate variants in genes associated with these disorders.

Whilst the ExAC dataset, with data from 60,706 population samples, represents a powerful resource for variant frequency filtering, some genomic regions, especially those that are repetitive or with high GC content are not fully covered. Specifically, 12.5% of sample bases across the 40 genes within CardioClassifier are covered at <20x (Supplementary Figure 3). In contrast, in our high-quality set of healthy volunteers 99.9% of sample bases are covered at >20x allowing us to accurately identify common and low-frequency variants or platform specific errors across, all regions of interest (rule BS1).

In addition to these in-house data, we also display counts from published clinical cohorts for HCM^9,20^, DCM^9,21^, LQTS^22^ and Brugada syndrome^23^.

### Results show high concordance with manually curated and gold-standard data

We compared CardioClassifier to 57 gold-standard, manually curated protein-altering variants from the recent ClinGen pilot for MYH7 (manuscript in submission, citation will be added). Of 222 rules activated by ClinGen for these 57 variants, 157 represented computationally accessible data (from 9 ACMG/AMP rules) that were fully retrieved by CardioClassifier. CardioClassifier concordantly activated 137/157 rules (87.3%; Figure 3). The discrepancies fall across 3 rules; PP3 (in silico prediction algorithms; n=12), PS4 (prevalence in affected individuals statistically increased over controls; n=7) and PM5 (same amino acid residue as known Pathogenic variant; n=1). CardioClassifier imposes a more stringent threshold on PP3 (allowing only one of eight in silico prediction algorithms to be discordant), and differences in PS4 and PM5 are due to the increased availability of proband data to the ClinGen team (not available from public repositories). In both cases, CardioClassifier successfully returned all available data.

**Figure 3:**
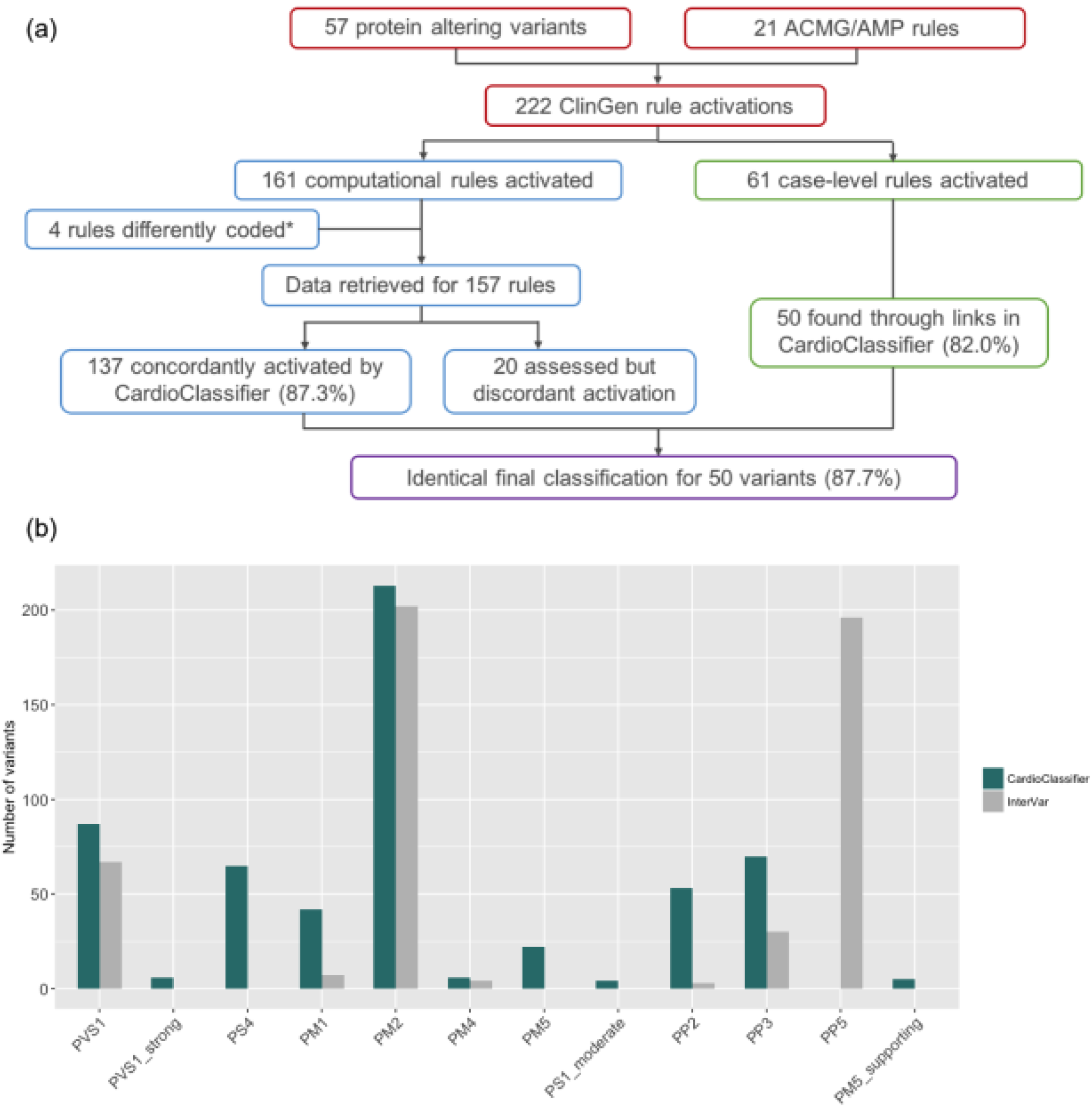
Validation of CardioClassifier. (a) Comparing CardioClassifier to 57 variant expertly curated by the ClinGen pilot for **MYH7**. Rules were split into those that can be computationally annotated and those that are ‘case-level’ and require manual input. CardioClassiifer was run using an ‘All Cardiomyopathy’ test to reflect the spectrum of phenotypes caused by variants in **MYH7**. *Of the computational rules, 3 were removed from the comparison as they represent draft modifications to the ACMG framework by the ClinGen Cardiovascular domain working group that are not yet published, and not yet implemented in CardioClassifier. Specifically, truncating variants in **MYH7** activate a new rule PVS1_moderate. Additionally, for variants classified as Benign by frequency alone (BA1) CardioClassifier does not assess any further rules, leading us to remove an additional data point from the comparison as we would not expect it to be retrieved. (b) Counts of individual rules activated by CardioClassifier and InterVar for 219 variants identified as Pathogenic or Likely Pathogenic in ClinVar. Only pathogenic evidence rules and rules activated by one of the tools at least once are shown.

We then tested the ability of the links within the CardioClassifier report to inform activation of the 61 case-level data points activated by the ClinGen team. These links allowed us to manually collate 50/61 (82.0%) individual data points with differences again in the availability of proband data (6 PS4_supporting, 1 PS4_moderate, 1 PS2, 1 BS4 and 2 PP1_moderate). After addition of this clinical data, we reached an identical classification to the ClinGen team for 50/57 (87.7%) variants (Figure 3a).

### CardioClassifier has higher sensitivity and specificity than non-specific interpretation tools

In February 2017 Li and Wang released InterVar, and its companion web-server winterVar, as the first tools that add automation into populating criteria from the ACMG/AMP guidelines^7^. Whilst these tools were crucial steps forward in application of the framework, they aim to support interpretation across the full spectrum of human genes and disorders.

To show the added value of the highly disease- and gene-specific annotations included in CardioClassifier, we compared CardioClassifier to InterVar using a set of 219 variants identified as ‘Pathogenic’ or ‘Likely Pathogenic’ on ClinVar, with high confidence, across six ICCs. Based on automatically-retrieved data only, InterVar identified 64/219 (29.2%) variants as Likely Pathogenic or Pathogenic, while CardioClassifier identified over double this number as clinically actionable (156/219) with a sensitivity of 71.2% (Supplementary Table 2). For both tools, sensitivity would be increased further through user addition of clinical and functional data.

Despite the lower sensitivity of InterVar, there are occasions where the tool activates rules inappropriately in the absence of gene-specific knowledge. Firstly, InterVar activates PVS1 in the TTN gene, regardless of protein location, when it is recognised that truncating variants in exons not constitutively expressed in the heart are not associated with DCM, and are commonly found in demonstrably healthy controls^15^. Consequently, InterVar will categorise rare variants in these regions as ‘Likely Pathogenic’ when in fact they are unlikely to be disease causing.

Secondly, InterVar activates rule PP5 (reputable source identifies the variant as Pathogenic) for 89.5% of the variants as they are reported as ‘Pathogenic’ in ClinVar.

The ACMG guidelines state that this rule should only be activated when the evidence supporting the classification is unavailable, yet this evidence is often contained within the appropriate ClinVar submission. Full details of the rules activated by both tools are shown in Figure 3b.

To ensure the higher sensitivity of CardioClassifier was not due to over-activating rules, we also tested a set of 67 ‘Benign’ and ‘Likely Benign’ variants from ClinVar across the same six ICCs. CardioClassifier identified 61/67 (91.0%) of these as Benign and the remaining 6 as VUS. Conversely, InterVar identified 41/67 (61.2%) as Benign with 22 as Likely Benign and 4 as VUS. Here InterVar activates BS2 when a variant is seen in the 1000 genomes dataset, which we believe is inappropriate for ICCs which do not fit the important caveat of ‘full penetrance expected at an early age’. We do acknowledge, however, that InterVar was developed for severe congenital and very-early onset developmental disorders with nearly 100% penetrance.

### Diagnostic yield in HCM cases matches previous reports

To investigate the clinical utility of CardioClassifier we used a dataset of 327 HCM cases. In 66 cases (20.2%) we immediately identified a Pathogenic (n=11) or Likely Pathogenic (n=55) variant, with a further 76 cases (23.2%) harbouring a VUS. To determine the proportion of these VUSs likely to become clinically actionable after the addition of case-level data, we calculated the excess of VUSs in cases over the background level of rare and presumably benign VUSs in 625 healthy volunteers (HVOLs). Based on a background level of 9.7%, we calculate a case excess of VUSs of 13.5%. Combining this with the 20.2% of cases with a Pathogenic or Likely Pathogenic variant, overall, 33.7% of cases have a potentially clinically relevant variant (Supplementary Figure 4a), comparable to previous reports^20^.

### Manual curation of known variants

In addition to automatic retrieval of computational data, CardioClassifier will store curated case-level data entered by users, or pre-populated from community curation efforts. We have primed this ‘knowledge base’ with data from 120 cardiomyopathy variants. We set out to create a ‘knowledge base’ of fully curated variants to include in CardioClassifier. This dataset comprised the 57 variants curated in the ClinGen MYH7 pilot study and the most commonly observed variants for the major cardiomyopathies; HCM, DCM and ARVC, defined as those that occurred six or more times in the ACGV resource (reflecting a HCM case frequency of ~1/1000)^9^. Collectively, the 84 variants in ACGV that were above this threshold are observed 1,258 times in 7,855 cardiomyopathy cases, representing 39.5% (1,258/3,186) of identified variants.

Of the 84 ACGV variants, 63 had not been curated by the MYH7 pilot. For these variants, we searched the literature and ClinVar for reports of segregation, de novo occurrence and functional characterisation. After this manual curation step, this dataset comprised 34 Pathogenic, 13 Likely Pathogenic and 7 variants that remain VUS (Supplementary Table 3; Supplementary Figure 4b). Addition of case-specific data did not alter the overall number of clinically actionable variants but increased the proportion of Pathogenic/Likely Pathogenic and downgraded a subset of variants to Likely Benign and Benign (3.9%). The annotations for these 120 variants have been stored in CardioClassifier to be retrieved and automatically included in classifications. Consequently, for at least 40% of variants identified in cardiomyopathy cases, CardioClassifier will immediately return the correct classification, with the remaining requiring some manual addition of case-level data where this has not been pre-loaded into the knowledge base.

## DISCUSSION

We describe CardioClassifier, an automated and interactive web-tool to aid clinical variant interpretation across a wide range of ICCs. To the best of our knowledge our tool represents a unique disease-specific solution that automates data retrieval, incorporates gene- and disease-specific knowledge to refine rule application, is pre loaded with curated data on prior observations (in health or disease), and integrates evidence according to the widely-adopted framework from ACMG/AMP. The tool is transparent, with all the information incorporated into interpreting each variant displayed along with the final classification. It is also flexible, and designed to be fully interactive, with the user able to add and remove evidence specific to the patient/family of interest.

The strength of CardioClassifier is its disease-specificity. The ACMG/AMP rules are intentionally non-specific to allow adoption in any disease domain. To harness the full power of this framework, the rules need to be applied in a disease- and gene-specific manner^24^. We have defined criteria and thresholds for each ACMG/AMP rule that are specific to the disorder of interest. We demonstrate the power and effectiveness of this approach through comparison with InterVar, a recently released genome-wide interpretation interface, which CardioClassifier significantly outperforms. Incorporation of disease-specific knowledge is, however, limited by our current understanding and the availability of disease datasets on which to perform analyses. Consequently, the power of this tool will continue to increase over time as new and informative data emerge across the full complement of ICCs. Furthermore, on-going community initiatives, such as the Clinical Genome Resource (ClinGen), are defining consensus disease and gene-specific standards for modifications to the ACMG/AMP guidelines. It is our intention to continue to develop CardioClassifier to utilise these standards as they become available.

We believe the main limitation to the effectiveness of any computational solution is the limited ability to retrieve all clinical or patient-specific data computationally. This is in part due to current databases, such as ClinVar, storing clinical assertions in free text fields, rather than as structured data that can be easily mined. The combination of CardioClassifier pre-populating computational data linked with the ability to interactively add additional levels of case and variant specific evidence in a structured format, will help to overcome this hurdle. Furthermore, we have manually curated and included case-level data pertaining to 120 variants in a variant specific ‘knowledge base’ and will continue to add to this as knowledge accumulates in the community. Future development of CardioClassifier will streamline data-sharing, to expand our knowledge base, and share this with the community via submission to the ClinVar database. This increasing knowledge base relies on researchers and clinicians in the field supporting data-sharing initiatives, and facilitating direct ClinVar submission from CardioClassifier for the benefit of the ICC community is a development priority. A further limitation to CardioClassifier in its current form is the restricted prediction of impact on splicing. This arises for two main reasons; firstly, CardioClassifier uses the Ensembl variant effect predictor to annotate variants, which annotates bases within 8 base-pairs of the exon/intron boundary as splice-site, but will miss more distal bone-fide splice site variants. Secondly, we currently have not incorporated any in silico splice-site prediction algorithms, due to limitations around availability, licensing and accuracy. These issues will be addressed in a future release.

CardioClassifier is designed to work seamlessly, but not exclusively, with the Illumina TruSight Cardio sequencing kit^1^, and takes as input the final output VCF from a standard bioinformatics analysis pipeline. We believe the combination of these two powerful tools is a crucial step in broadening the availability of genetic testing for ICCs, and standardising variant interpretation in this field. Furthermore, we hope that in demonstrating the clinical utility of our disease-specific approach, we will encourage others to develop similar tools across other disease specialties.

## TOOL AVAILABILITY

CardioClassifier is available at www.cardioclassifier.org, with a free license for non commercial use. The code and data used to produce this manuscript will be available at: https://github.com/ImperialCardioGenetics/CardioClassifierManuscript.

## ACKNOWLEDGEMENTS

This work was supported by the Wellcome Trust (107469/Z/15/Z), the Medical Research Council (UK), the Cardiovascular Research Centre Biobank at Royal Brompton and Harefield NHS Foundation Trust (NRES ethics number 09/H0504/104+5), the Fondation Leducq (11 CVD-01) and a Health Innovation Challenge Fund award from the Wellcome Trust and Department of Health, UK (HICF-R6-373).

This publication includes independent research commissioned by the Health Innovation Challenge Fund (HICF), a parallel funding partnership between the Department of Health and Wellcome Trust. The views expressed in this work are those of the authors and not necessarily those of the Department of Health or Wellcome Trust.

## SUPPLEMENTARY INFORMATION

- SUPPLEMENTARY NOTE 1 - Parameterisation of ACMG/AMP rules
- SUPPLEMENTARY NOTE 2 - Benchmarking against manually curated variants
- Supplementary Table S1
- Supplementary Table S2
- Supplementary Table S3
- Supplementary Figure S1
- Supplementary Figure S1
- Supplementary Figure S2
- Supplementary Figure S3
- Supplementary Figure S4

